# Architectural underpinnings of stochastic intergenerational homeostasis

**DOI:** 10.1101/2023.11.15.567256

**Authors:** Kunaal Joshi, Charles S. Wright, Rudro R. Biswas, Srividya Iyer-Biswas

## Abstract

Living systems are naturally complex and adaptive, and offer unique insights into the strategies for achieving and sustaining stochastic homeostasis in different conditions. Here, we focus on homeostasis in the context of stochastic growth and division of individual bacterial cells. We take advantage of high-precision longterm dynamical data that have recently been used to extract emergent simplicities and to articulate empirical intra- and intergenerational scaling laws governing these stochastic dynamics. We identify the core motif in the mechanistic coupling between division and growth, which naturally yields these precise rules, thus also bridging the intra- and intergenerational phenomenologies. By developing and utilizing novel techniques for solving a broad class of first passage processes, we derive the exact analytic necessary and sufficient condition for sustaining stochastic intergenerational cell size homeostasis within this framework. Furthermore, we provide predictions for the precision kinematics of cell size homeostasis, and the shape of the interdivision time distribution, which are compellingly borne out by the high-precision data. Taken together, these results provide insights into the functional architecture of control systems that yield robust yet flexible stochastic homeostasis.

---

Robust architecture is a common feature of functional complex and adaptive systems. Strict constraints on protocols enable a plug-and-play modularity that confers flexibility to (or “deconstrains”) the overall systems design (1, 2). Recent high-precision experiments and analysis of extant data on different microorganisms have shown that stochastic intergenerational homeostasis of cell sizes is constrained by surprisingly universal and elegant emergent simplicities (3, 4), despite the substantial differences in underlying molecular circuitry governing growth and division in system- and environment-specific ways. What robust architectures lead to the observed intra- and intergenerational emergent simplicities governing stochastic intergenerational homeostasis?

Between successive divisions, cell size increases stochastically while adhering to an *intra*generational scaling law: the mean-rescaled cell size distributions of cells at different times since the last division event undergo a scaling collapse (5, 6) (see Fig. 1c). The mean itself increases exponentially with time since the last division event (5, 6). Furthermore, *inter*generational size dynamics are Markovian and a scaling law constrains the precision kinematics of stochastic intergenerational homeostasis: the mean-rescaled size-at-birth in the next generation distributions are independent of the sizes-at-birth in the current generation (3, 4) (see Fig. 1d). Intuitively it is clear that these empirically observed scaling laws or “emergent simplicities” must reflect key aspects of the nature of the coupling of growth to division, but using the observed phenomenology to decipher the underlying mechanism has remained an open challenge. Here we provide the solution to this problem.

**Fig. 1.**
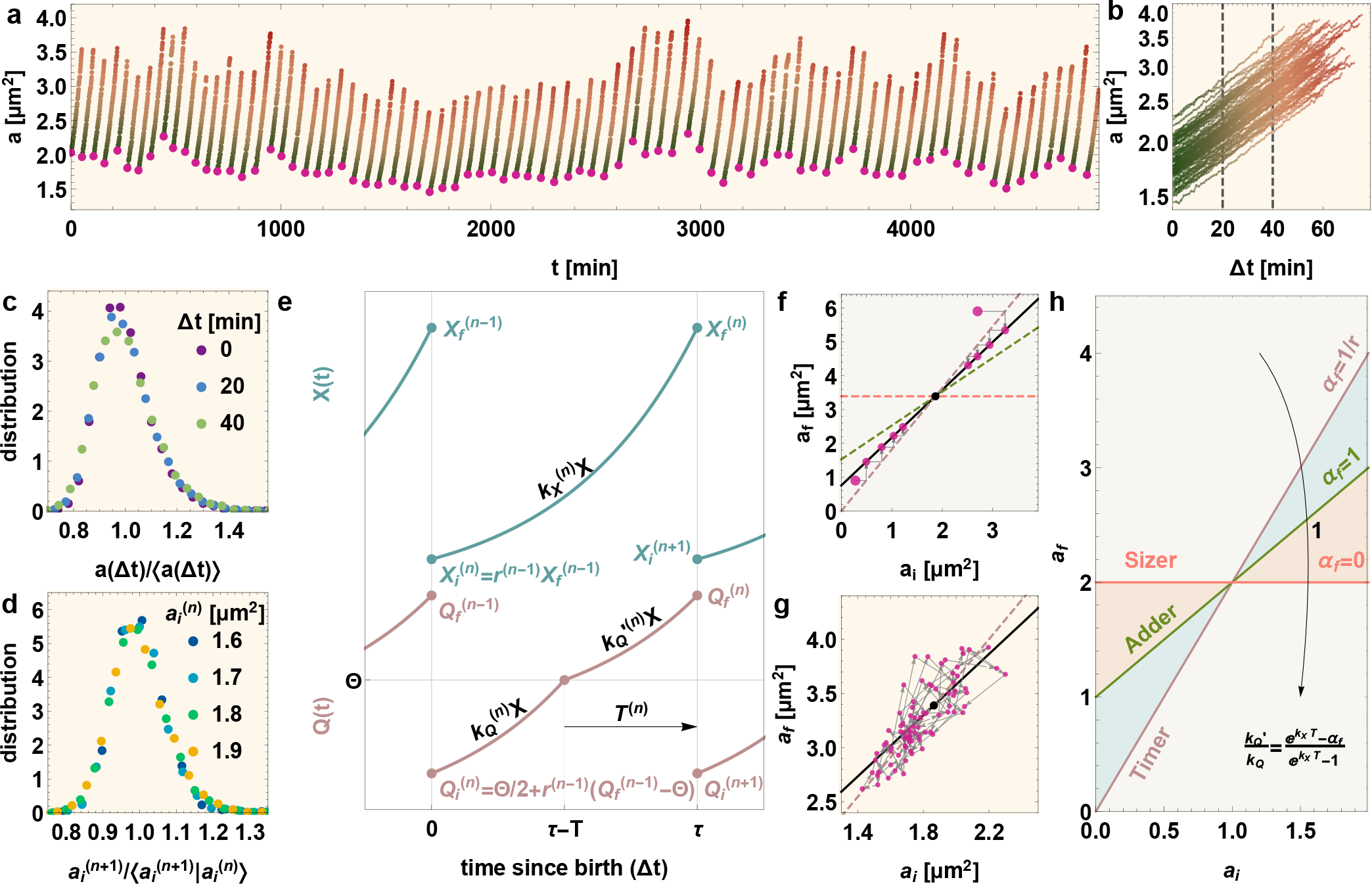
Empirically observed emergent simplicities motivate the mechanistic model for stochastic intergenerational homeostasis. (**a**) Stochastic intergenerational homeostasis of cell sizes-at-birth (highlighted magenta circles) as seen in high-precision data recording an individual cell’s stochastic growth and division dynamics over multiple generations. **(b)** The cell sizes in (a), are replotted on a log-linear scale versus time since the last division event (Δ*t*); cell sizes undergo stochastic exponential growth between divisions. **(c)** The distributions of cell sizes at different times since birth (marked by the dashed gray lines in (b)) are plotted after rescaling by their respective mean values. These mean-rescaled distributions undergo a scaling collapse, an intragenerational scaling law consistent with the Stochastic Hinshelwood Cycle model of stochastic exponential growth. **(d)** The conditional distributions of next generation’s initial sizes given the current generation’s initial size are plotted for different current initial sizes, after rescaling by their corresponding mean values. These mean-rescaled distributions undergo a scaling collapse revealing an intergenerational scaling law, which in turn specifies the precision kinematics of stochastic cell size homeostasis. **(e)** The proposed mechanistic model bridges the intra-generational and inter-generational cell growth and division dynamics. *X* is the effective Hinshelwood cycle variable corresponding to cell size, while *Q* represents the thresholding protein. 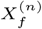 and 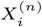 are the copy numbers of *X* and *Q* at birth in the *n*^*th*^ generation, and 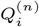 and 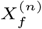 are the copy numbers at division. Throughout the cell cycle, *X* is produced at rate 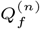. Initially, *Q* is produced at rate 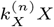 until it crosses the threshold at Θ, after which time its production rate is reset to 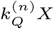. After crossing the threshold, cell division occurs after time *T* . Upon division, the next generation’s *X*_*i*_ and *Q*_*i*_ values are related to the current generation’s *X*_*f*_ and *Q*_*f*_ values through the division rules given by Eq. (10). The proposed *stochastic* model naturally yields the observed phenomenologies in (a), (b), (c), (d), and (g). **(f)** Heuristic argument for intergenerational homeostasis in extant models based on the deterministic sizer–timer–adder paradigms. Within this scheme, the inter-generational final size (*a*_*f*_) vs initial size (*a*_*i*_) dynamic is thought to occur as shown: starting from an initial generation characterized by the coordinates of the large red dot, the cell deterministically adjusts its size to exponentially relax to the target cell size set by the black point, the intersection between lines corresponding to the growth and division rules. **(g)** In contrast to the heuristics suggested by the extant sizer–timer–adder paradigms (shown in (f)), the experimentally observed high-precision Inter-generational *a*_*f*_ vs *a*_*i*_ trajectories (here taken from the cell shown in (a)) are dramatically and quantitatively different, thus motivating the necessity for a completely revised framework. **(h)** In the appropriate ranges of parameter values, the fully stochastic mechanistic model we propose here can recapitulate specific mean behaviors displayed by the sizer–timer–adder paradigm. For the mean of final size given initial size on initial size in the quasi-deterministic limit, the slope (*–*_*f*_) is controlled by the relative rate of production of *Q* after crossing threshold 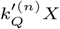. In the deterministic limit, slopes *–*_*f*_ = 0, 1, 1*/r* (with 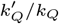 greater than, equal to, and less than *k*_*Q*_ respectively) correspond to sizer, adder, and timer models, where *r* is the deterministic division

In addition to yielding the observed emergent simplicities, the minimal mechanistic model we propose here has inbuilt constraints that deconstrain, allowing for versatile implementations with different system-specific details for different microorganisms (or even growth conditions), while robustly ensuring that homeostasis will result in each instantiation despite the inherent stochasticity in the growth and division processes. From the point of view of evolvability, conserved core functional architectures serve to constrain variation that would break the core mechanism. On the balance, they confer flexibility and robustness to processes that leave the core intact (1, 2).

We start with the minimal model that reproduces the observed universal statistics of cell size growth, namely the Stochastic Hinshelwood Cycle (SHC) model of stochastic exponential growth (5–8). Let *X* represent the effective SHC variable undergoing stochastic exponential growth according to (5, 8):

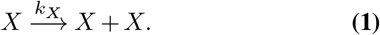

It relates to the cell size, *a*, via:

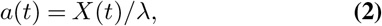

where λ is a scaling factor relating the discrete copy numbers of *X* to the cell size, *a*. The previously noted intragenerational scaling law is consistent with this model, since in balanced growth conditions it naturally yields:

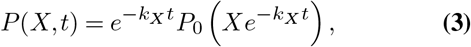

where *P*_0_ is an initial condition-dependent distribution (5– 8). Since the mean grows as 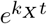 with time, when this distribution at any given time is rescaled by its mean value, a time-invariant distribution results.

Additionally, since the cell size-at-birth distribution must satisfy the intergenerational scaling governing stochastic intergenerational homeostasis, so must the copy numbers of *X* at “birth”, i.e., immediately following a division event (3, 4):

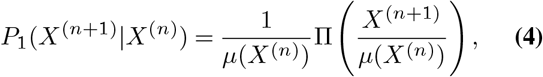

where *P*_1_ is the conditional distribution of *X*^(*n*+1)^ (the next generation’s initial copy numbers, i.e., copy numbers at birth) given *X*^(*n*)^ (the current generation’s initial copy numbers). *μ* is the next generation’s mean initial copy number as a function of the current generation’s initial copy number, and Π is the invariant distribution that results after mean-rescaling *P*_1_. This emergent simplicity, as we have derived in (4), specifies the precision kinematics of initial copy numbers over successive generations through the exact stochastic map:

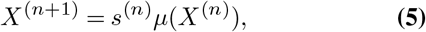

where 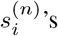 are random numbers drawn from the distribution II (with unit mean); the superscript serves to record the generation as *n*. In sum, in the formulation we have presented here, the specific challenge is to bridge intra- and intergenerational phenomenologies by identifying the correct mechanistic coupling between growth and division that naturally yields the intergenerational scaling law, Eq. (4).

## Results

### (I) Mechanistic underpinnings of stochastic intergenerational homeostasis

A minimal model consistent with empirical observations can be articulated as follows (see Fig. 1 for a graphical summary). As outlined in Eqs. (1) and (2), the copy numbers of *X* serve as a proxy for cell size *a* and undergo stochastic exponential growth. The mechanism of size control is implemented by an auxiliary “growth reporter”, *Q*, whose numbers increase stochastically with a propensity proportional to the copy numbers of *X* present:

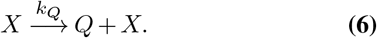

When *Q* reaches a threshold value of Θ, the decision to commit to division is taken and a stochastic process commences culminating in cell division after a random delay time *T* . In this post-threshold period (of duration *T*), the *X*-*Q* dynamics continue to proceed as in Eqs. (1) and (6); however the propensity of production of *Q* may differ and is thus denoted by 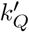. Finally, cell division occurs with the copy numbers reset according to Eq. (10) and governed by the division ratio *r*, a random variable we assume to be independent of cell size. For symmetrically dividing cells, the mean division ratio is 1*/*2.

### (II) Intragenerational statistics: Exact analytic solution

While several techniques are known for solving for stochasticity arising due to copy number fluctuations in different models (9–14), an exact analytic solution to coupled stochastic evolution of *X* and *Q*, as encoded in Eqs. (1) and (6) respectively, is not readily derived via traditional approaches. Instead, we solve this seemingly intractable problem (below) through a “stochastic rescaling” of time. Our method relies on the fact that while *X* influences the growth of *Q* through the rate *k*_*Q*_ *×X, Q* does not influence the stochastic growth dynamics of *X*. Our mathematical technique is broadly applicable to scenarios where the growth rates for both *Q* and *X* are arbitrary functions of *X*.

We define a new rescaled time variable, *t*_*r*_, whose rate of change with the laboratory time variable, *t*, is just the growth rate of *Q*:

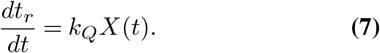

When the laboratory time *t* is replaced by *t*_*r*_, from Eqs.(1) and (6) we see that the dynamics of *Q* becomes formally *X*-independent, while the growth rate of *X* becomes the ratio of its laboratory growth propensity to that of *Q*, also formally independent of *X*! Thus, when the time variable is *t*_*r*_, the dynamics become that of two uncoupled growth reactions, schematically represented thus:

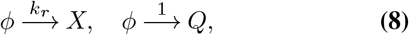

with *k*_*r*_ = *k*_*X*_ */k*_*Q*_. In terms of this rescaled time, the coevolution of *X* and *Q* can be obtained analytically, even though characterizing their co-evolution in laboratory time is difficult. Specifically, using standard techniques of stochastic processes (9, 10), we have calculated analytically that the distribution of *X* = *X*_Θ_ when *Q* reaches the threshold value of Θ, when starting from initial values (*X*_*i*_, *Q*_*i*_), is a Pascal distribution:

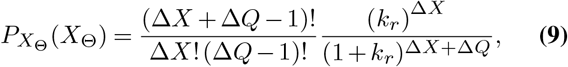

where Δ*X* = *X*_Θ_ − *X*_*i*_ ≥0 and Δ*Q* = Π − *Q*_*i*_ ≥1 (see Appendix Sec. A).

#### The quasi-deterministic limit

We now consider an interesting limit of this process, which is useful for comparison with experimental data. From the above distribution, the ratio of the standard deviation to the mean for Δ*X* is 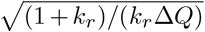. Since *k*_*r*_≫1 is the physical regime of interest where the numbers of *Q* are much smaller compared with numbers of *X*, and since in steady state Δ*Q*∼ Θ up to a fractional factor 1*/*2, we find that for large Θ the standard deviation becomes negligible compared to the mean (their ratio becomes 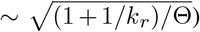. In this regime, the distribution of Δ*X*_Θ_ *≃ k*_*r*_Δ*Q* becomes an almost deterministic function of *Q*_*i*_. Furthermore, if 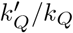 is negligible and division noise is limited, *Q*_*i*_ is just a constant times Θ. In summary, for Θ *∫* 1+ 1*/k*_*r*_, .6.*X*_8_ becomes quasi-deterministic; furthermore, when 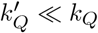, its value is a function only of 8 and *k*_*r*_ and thus independent of *X*_*i*_ (see Appendix Sec. D). In other words, in this limit, a constant amount is added to *X*_*i*_ during the time taken for *Q* to reach the threshold.

As outlined previously, once the thresholding of *X*_Θ_ occurs, a division process commences that takes time *T*, following which the cell divides with a division ratio *r*. (Note that both *T* and *r* are random variables whose values change from generation to generation.) During this process, *X* and *Q* continue to grow following Eqs. (1) and (6), with 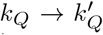 . We have analytically solved the corresponding Master Equation and found the joint moment-generating function for the final pre-division copy numbers, (*X*_*f*_, *Q*_*f*_), starting from the respective values at the threshold, (*X*_Θ_, Θ) (Appendix Sec. B). Combining the analytic results (Eq. (9)) for the statistics of *X*_Θ_, we have analytically calculated the statistics of (*X*_*f*_, *Q*_*f*_), given initial values (*X*_*i*_, *Q*_*i*_) (see Appendix Sec. B). These statistics completely specify the *intra*generational stochastic evolution of cell size in our framework and are used in the following sections.

### (III) Intergenerational statistics: Homeostasis condition

We now proceed to determine the *inter*generational evolution of (*X, Q*), and hence the cell size. This is provided by the division rule that converts the final pre-division values in a given generation, (*X*_*f*_, *Q*_*f*_), to the initial values, (*X*_*i*_, *Q*_*i*_), in the next generation. Using the notation *A*^(*n*)^ to represent the value of a random quantity *A* measured in generation *n*, we propose the following division rules that incorporate the cell size division ratio *r*:

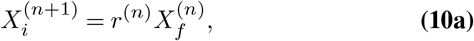

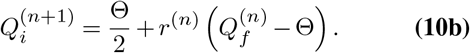

For asymmetrically dividing cells, we underscore the subtle point that a portion of *Q*, equal to the threshold amount Θ, is split equally during division among the daughter cells, and not at the division ratio *r*. Biochemically, such behavior may naturally arise, for instance, if Θ amount of *Q* accumulates around the cell division plane to initiate division. This assumption is not necessary for achieving cell size homeostasis but is consistent with the experimentally observed simplicities discussed in the following sections. The implications of alternate division rules are explored in Appendix Sec. E. In a later section, we discuss possible biological implementations and implications in greater detail.

We can now consider the question of the homeostatic stability of the intergenerational evolution of *X* and consequently cell size. Addressing this problem requires consideration of the intergenerational co-evolution of *both X* and *Q*. Yet, since the absolute amount of *Q* is constrained at the thresholding point in every generation, *Q* is trivially in homeostasis. We can thus simply consider the intergenerational evolution of the value of *X* at a fixed point in the cell cycle. Specifically, we choose to follow the intragenerational evolution of *X*_Θ_, since at that thresholded event the value of *Q* must be Θ and so its co-evolution is trivial. We find the following intergenerational evolution of the reaction noise-averaged moments of *X*_Θ_, *μ*_*m*_ = ⟨(*X*_Θ_)^*m*^⟩ for *m≥* 1 (see Appendix Sec. B for details; define *μ*_0_ = 1):

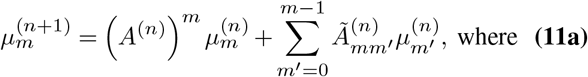

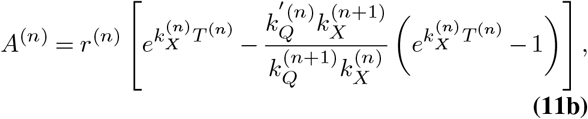

and 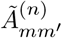 are bounded quantities whose exact forms are unimportant for the homeostasis of *X*_Θ_ and thus, cell size. Under intergenerational evolution in accordance with the above equations, attainment of stochastic homeostasis is assured, provided that all moments 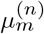, as *n*→ *∞*, (i) become independent of the initial value 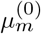, and (ii) remain finite. Assuming that the *A*^(*n*)^ are uncorrelated for different *n*, being independent draws of a random variable *A*, we have derived (see Appendix Sec. C) the necessary and sufficient conditions for cell size homeostasis as the following set of bounds on the moments of *A*:

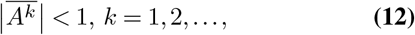

where the overline denotes an average over generations. As shown previously in (4), this sequence of conditions is equivalent to a simple bound on *A*: |*A*| _max_ ≤1. (The inequality is strict only if *A* is deterministic. If the inequality is violated for *k* = *k*_0_, all moments *μ*_*k*_ with *k*≥ *k*_0_ are unstable, i.e., do not reach homeostatic initial-condition-independent finite steady-state values (4).) These general conditions for homeostasis in our model are reminiscent of conditions derived in the phenomenological theory in (4), corresponding to emergent simplicities in cell size homeostasis observed in experiments (3).

#### The quasi-deterministic limit

The quasi-deterministic limit applies when the copy numbers of *X* and *Q* are large enough that the reactions in Eqs. (1) and (6) proceed deterministically. This applies when *X*_*i*_, *Q*_*i*_ *≫*1 (consistent with the condition Θ ≫1+ 1*/k*_*r*_ considered earlier). In this limit, intragenerational dynamics proceed deterministically, and the primary source of noise in the system is due to the intergenerational variation of reaction rates 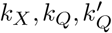 and the duration between threshold-crossing and division, *T* . By Eq. (1) our model cell undergoes quasi-deterministic exponential growth, in agreement with high-precision experimental observations of exponential growth in bacterial cells under constant nutrient-rich growth conditions (6). Meanwhile, intergenerational evolution of cell size is encapsulated in the relation between initial sizes of successive generations (see Appendix Sec. D):

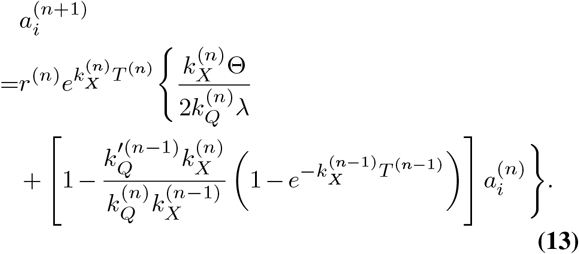

#### Recapitulating known results for mean behaviors

The mean of final size (*a*_*f*_) given initial size (*a*_*i*_) is found to vary linearly with the initial size for nearly all bacterial species studied (15):

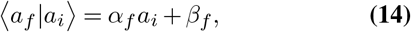

for constants *α*_*f*_ and *β*_*f*_ that are species- and condition-dependent. Traditional deterministic homeostasis models consider the final size *a*_*f*_ of a cell with initial size *a*_*i*_ to be equal to ⟨*a*_*f*_ |*a*_*i*_⟩, and hence to follow the deterministic map 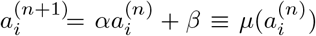, where *α* and *β* are equal to ⟨*r*⟩ *α* _*f*_ and ⟨*r* ⟩*β*_*f*_, respectively, ⟨*r*⟩ is the average division ratio and *μ* is the mean function in Eq. (5). For such a deterministic map, *a*_*n*_ converges to a finite value independent of *a*_0_ as *n → ∞*iff | *α*| *<* 1 (16, 17). This formulation has been used for the adder (*α* =⟨ *r*⟩), timer (*α* = 1), and sizer (*α* = 0) models (18–23) (Fig. 1 f, h). Although these models adequately describe mean trends, they fundamentally fail to capture the observed stochastic dynamics, governed by the stochastic map given by Eq. (5),

which results from the intergenerational scaling law, Eq. (4). That the mythical ‘average’ cell fails to capture the stochastic behaviors of the individual bacterial cell is increasingly well appreciated in different contexts (3, 4, 24–29). Starting with Eq. (13), taking an intergenerational average over the stochastic variables 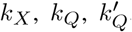, and *T* results in the prediction that the conditional mean of the next generation’s initial size given the current generation’s initial size varies linearly with the current generation’s initial size, consistent with observations above. Moreover, the ratio 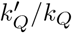 can be used to tune the slope *α* (Fig. 1h). Slopes between pure adder and pure timer require 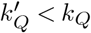, the slope for the pure adder requires 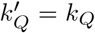 (or trivially when *T* = 0 and the model becomes 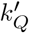 -independent), and slopes between pure sizer and pure adder can be obtained when 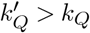.

### (IV) Comparison with data: emergent simplicities and parameter extraction

Incorporating into Eq. (5) the observed linear dependence of the mean, *μ*(*a*)= *α a* + *β*, which is also reproduced by our model (see preceding section), the significant emergent simplicity governing intergenerational cell size evolution is (3, 4):

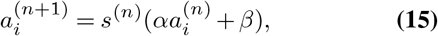

where 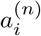 is the initial newborn size in the *n*^th^ generation; the numbers *{s* ^(*n*)^ *}* are independent random instantiations of a random variable *s* with unit mean and a growth conditiondependent probability distribution; and (*α, β*) are the growth condition-dependent constants determining the mean *μ* (Fig. 2a).

**Fig. 2.**
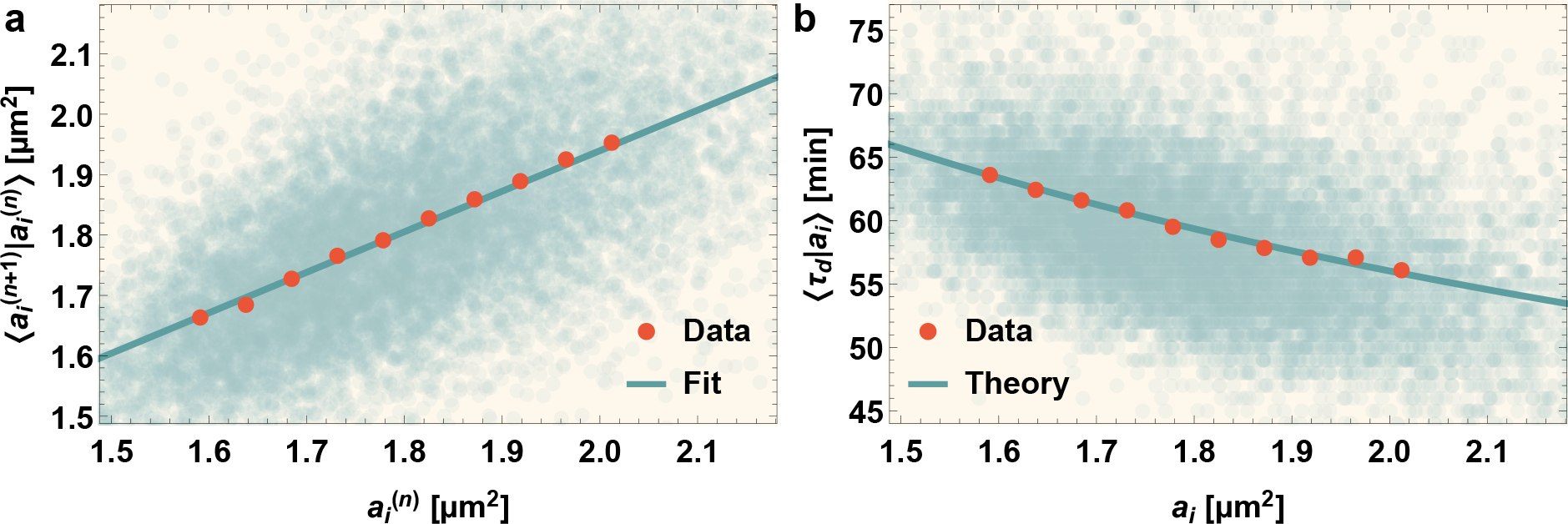
Parameter estimation; theoretical predictions are validated by experimental data. (**a**) The experimentally measured mean (red, with error bars) of next generation’s initial area is plotted as a function of current generation’s initial area. The only fitting parameter in the model, *k*_*r*_ Θ */λ*, is estimated as twice the intercept over slope of the linear fit to mean (see Eq. (17)). The light blue scatter plot in the background shows next generation’s initial area vs current initial area for different cell cycles in the data. **(b)** The experimentally measured mean (red, with error bars) of division time is plotted as a function of initial area. The blue curve is the analytic model prediction given by Eq. (23). Note that the error bars are small compared to marker size and hence not visible.

This simplicity straightforwardly emerges from our model in the quasi-deterministic limit when 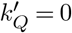 and Θ ≫ 1+ 1*/k*_*r*_ (also, *X* ≫*Q*). Here, we can set 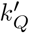 to zero since the experimental data showing this emergent simplicity is obtained from *Caulobacter crescentus* cells: for these cells, the slope of the conditional mean of the next generation’s initial size given the current generation’s initial size lies between those of the pure adder and pure timer, hence 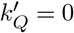 satisfies the required constraint 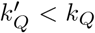. Since 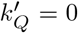 at the end of each cell generation *Q* = *Q*_*f*_ = Θ. Using Eq. (10), in steady state the initial amount of *Q* is always *Q*_*i*_ = Θ */*2. As observed previously, when Θ ≫1+ 1*/k*_*r*_, *X*_Θ_ is quasi-deterministic and results from adding a constant amount to *X*_*i*_: *X*_Θ_ ≃*X*_*i*_ + *k*_*r*_Δ*Q* = *X*_*i*_ + *k*_*r*_Θ*/*2. Due to quasi-deterministic exponential growth through the period *T* of the subsequent division process, *X* increases further to 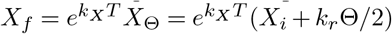). Converting from *X* to cell size *a* using Eq. (1) and applying Eq. (10), our model yields:

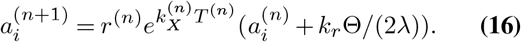

Taking into account the intergenerational stochasticity of *r, T* and *k*_*X*_, this stochastic map is equivalent to the emergent scaling law for intergenerational cell size control, Eq. (15), obtained from experimental data! We can identify the observed constants in this law with parameters of our model:

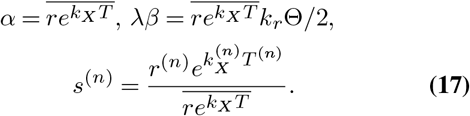

As before, the overline denotes an intergenerational average and *k*_*r*_ = *k*_*X*_ */k*_*Q*_ (assumed constant). Conversely, we can estimate the following model parameters and distributions from intergenerational growth and division data (see Fig. 2a):

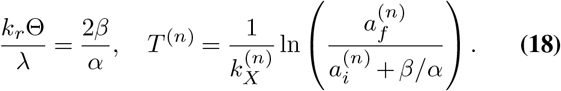

Note that the first relation in the above equation applies not only in the quasi-deterministic limit, but also in the non-approximate case. We have extracted the values of *α* and *β* from experimental data in Fig. 2a and shown predictions match consistent with Eq. (14) in Fig. 2b.

In conclusion, *k*_*X*_ is the experimentally measured cell size growth rate, 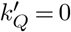, and *k*_*Q*_ is proportional to *k*_*X*_ with a constant of proportionality *k*_*r*_ = *k*_*X*_ */k*_*Q*_. The constant *k*_*r*_Θ*/λ* is the only fitted parameter in our model, obtained through Eq. (18) and the extracted values of *α* and *β* from the data fit in Fig. 2a. Once this is obtained, *T* can be measured from individual cell cycles through Eq. (18), and the joint distributions of *r, T* and *k*_*X*_ compiled. The data do not yield values of Θ, *k*_*r*_ (equivalently, *k*_*Q*_ = *k*_*X*_ */k*_*r*_) or *λ* individually, however these are constrained by our assumption of intragenerational noise-free growth (Θ ≫1+ 1*/k*_*r*_) and allow for a range of combinations that provide data–theory matches. Combining with Eq. (18), we require for self-consistency:

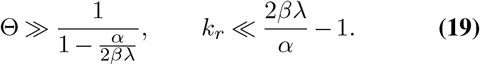

For large values of *β λ*, which correspond to large numbers of *X* in the cell, the first condition becomes simply Θ ≫1, i.e., the cell contains a large number of *Q*, even though these may be far fewer in number than *X*.

### (V)Exact solution and robust predictions

With 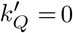, the exact solution for the distribution of copy numbers of *X* at division (*X*_*f*_) given initial copy numbers (*X*_*i*_) is (see Appendix Sec. F),

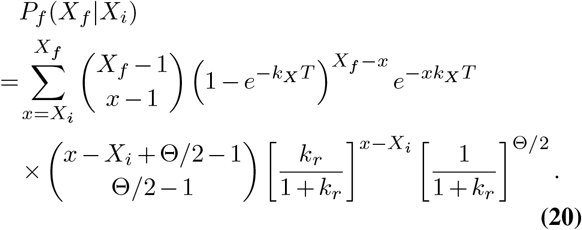

The above distribution is the predicted distribution for a single cell cycle with given *k*_*X*_ and *T* values. The overall distribution can be obtained by taking the intergenerational average with respect to the (observed) joint distribution of *k*_*X*_ and *T* values. From this analytic result we can find the distribution of the next generation’s initial size by multiplying the current generation’s final size (equal to *X*_*f*_ */λ*) by the division ratio (*r*), then taking the intergenerational average with respect to the observed joint distribution of *k*_*X*_, *T*, and *r* values.

Our analytic results for the distribution of the next generation’s initial cell size, conditioned on the current generation’s initial cell size, are compared with experimental data in Fig. 3. There is superb agreement between experiment and theory. The exact size distributions predicted by our model also undergo the experimentally observed intergenerational scaling collapse. Our mechanistic model can thus generate the experimentally observed multigenerational size data on single cell growth and division with quantitative accuracy. Furthermore, we reiterate that our model predictions robustly match these dynamics irrespective of the exact choice of model parameters, provided the chosen parameters satisfy the constraints given by Eqs. (18) and (19).

**Fig. 3.**
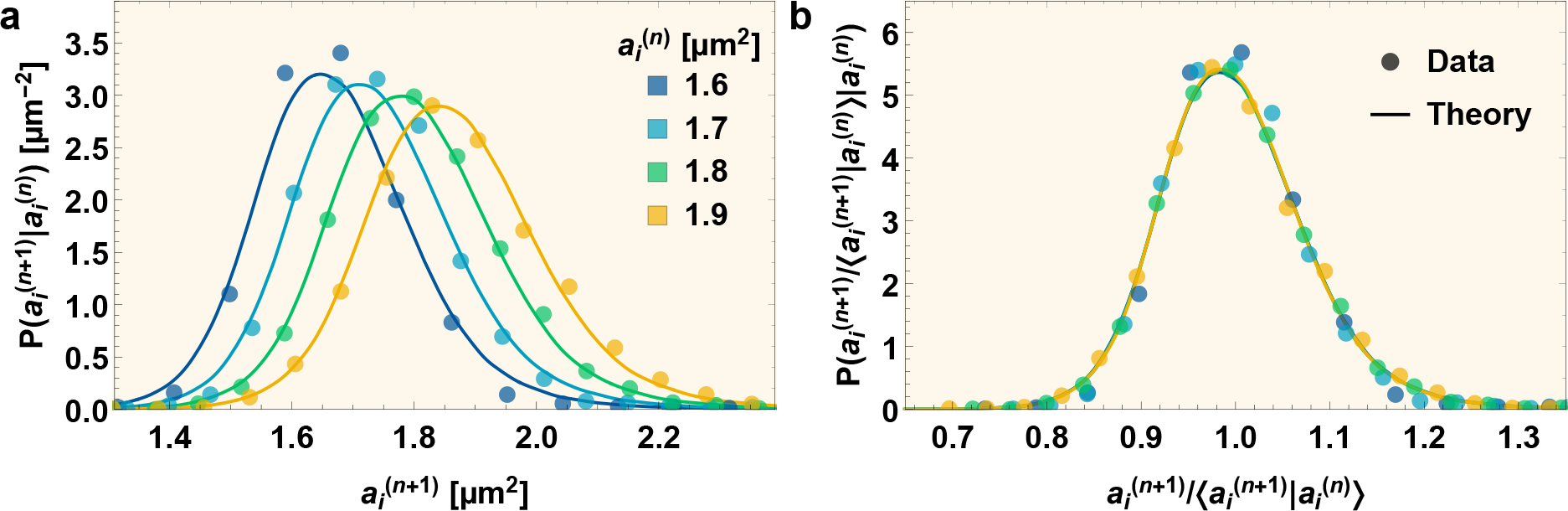
Intergenerational scaling law: experimental data and predictions from the mechanistic model. (**a**) Conditional distributions of next generation’s initial areas 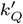 given current initial areas 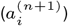, plotted for four different current initial areas (marked by different colors). The solid lines are the results of exact model simulations while the points represent experimental data. **(b)** Both experimentally measured and theoretically calculated distributions in (a) overlap when rescaled by their respective mean values.

### Condition for stochastic intergenerational size homeostasis

Given the scaling rule Eq. (15) for the intergenerational evolution of cell size, the conditions governing cell size homeostasis have been shown to be (4):

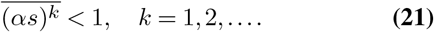

Since obtaining Eq. (15) from our model necessitated setting 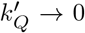, using this condition in Eqs. (17) and (11b) we find that *α s* = *A*. Thus the experimentally relevant cell size homeostasis conditions, Eq. (21), are identical to the more general conditions corresponding to homeostasis in our model, Eq. (12), in the limit where our model is consistent with experimental data!

In (3), we have shown that the stochastic map Eq. (15) accurately predicts the observed dynamics of initial cell sizes over successive generations, leading to cell size homeostasis. However, we have shown above that this stochastic map is also obtained in the quasi-deterministic limit of our mechanistic model, which should enable us to generate the full intergenerational evolution of cell sizes. In Fig. 4, we show this evolution starting from different initial sizes. Our model accurately predicts the observed distributions of cell sizes over successive generations leading to cell size homeostasis.

**Fig. 4.**
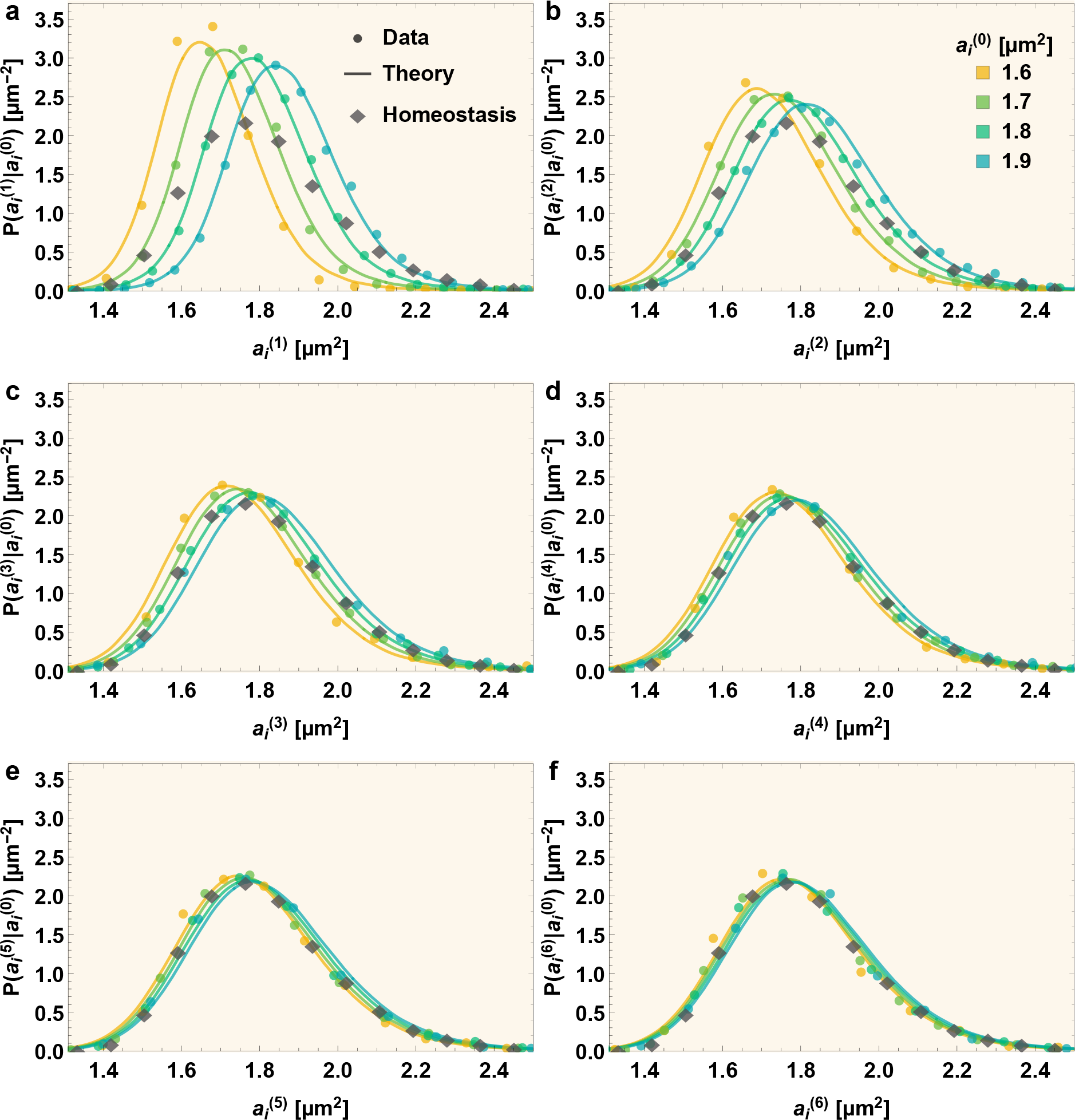
Precision kinematics of stochastic intergenerational homeostasis: experimental data and predictions from the mechanistic model. The conditional distributions of initial sizes after *n* generations 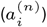, conditioned on the initial generation’s initial size 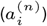, are plotted in (a-f) for *n* =1 to 6 respectively. The four different starting initial areas 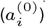 correspond to different colors. The solid lines are theoretical predictions based on exact simulations of the mechanistic model, while the points are experimentally measured data. The diamond markers denote the experimentally measured population wide homeostatic initial area distribution. All conditional distributions converge to this distribution as *n* increases, irrespective of the starting initial area.

### The division time distribution

Using the quasi-deterministic limit prediction Eq. (18), the cell division time, *τ* = ln(*a*_*f*_ */a*_*i*_)*/k*_*X*_, becomes:

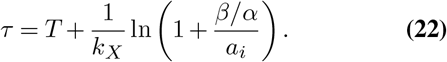

Here, *τ*, *T*, *k*_*X*_, and *a*_*i*_ are from the same generation. From this, we predict that when *a*_*i*_ is fixed, the mean division time is just:

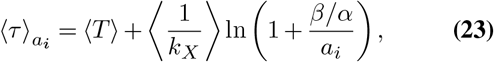

where ⟨ · ⟩and 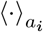 denote averaging over all generations or generations restricted by initial size value *a*_*i*_, respectively. This predicted functional form is compared against experimental values of 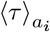 in Fig. 2c, showing excellent agreement.

Furthermore, using the experimentally measured joint distribution of *k*_*X*_ and *T*, 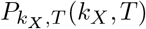, and the modelpredicted steady-state initial size distribution, 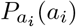, obtained through numerical methods described above and shown as the theoretical initial condition-independent homeostatic distribution in Fig. 4 f, our framework yields both the detailed conditional division time distribution for a given initial size, *P*_*τ*_ (*τ* |*a*_*i*_), and, by averaging over *a*_*i*_ using the homeostatic size distribution, the full steady-state division time distribution, *P*_*τ*,*ss*_(*τ*):

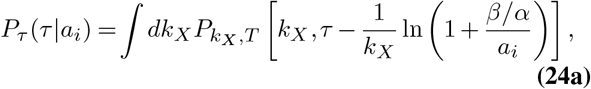

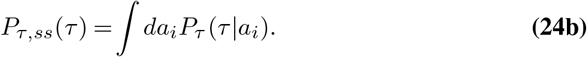

In Fig. 5 we show both conditional and full steady state division time distributions obtained through the *exact* Gillespie simulations of our mechanistic model. While the chosen model parameters (same as used to derive analytic results in previous sections) satisfy the conditions for the quasideterministic limit, the simulations are exact and do not assume quasi-deterministic simplifications. Once again, predictions match compellingly with the corresponding experimental data (see Fig. 5).

**Fig. 5.**
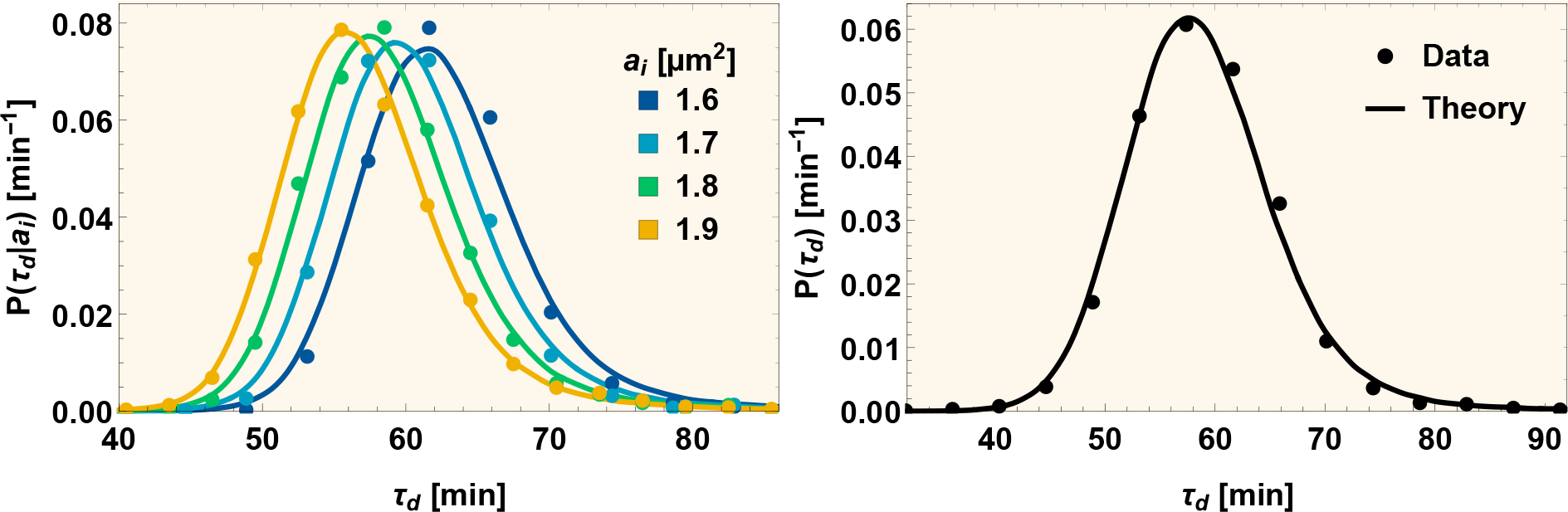
Shape of the interdivision time distribution: experimental data and predictions from the mechanistic model. **(a)** Division time distributions disambiguated by initial area are plotted for different initial areas (distinguished by different colors). The solid lines are the theoretical predictions from the mechanistic model, while the points are experimentally measured data. **(b)** The full division time distribution.

### Biological identities of X and Q

Bacterial cell division involves assembly of the division machinery (divisome) followed by cell wall constriction and ultimate cleavage (30). One of the earliest models of cell division hypothesized a diffusible factor that initiates division upon accumulation to a critical level (31); this factor was later suggested to be the tubulin homolog FtsZ, whose assembly dynamics are driven by cell growth rate (32). FtsZ is a critical player in recruiting and regulating members of the divisome, including cell wall remodelers responsible for synthesis and placement of peptidoglycan (PG) at the site of constriction (33, 34), via formation of the the Z-ring at the future division site (35). FtsZ exists in two conformations, found in monomer form or in (proto)filaments, respectively (36), which exhibit cooperative assembly such that additional monomers above a critical concentration increase only the polymer concentration (37, 38). The structure of the Z-ring is dynamic, with FtsZ exhibiting “treadmilling” (continuous polymerization and depolymerization at opposite ends of a filament) (39), typically at a rate of 30–40 nm/s, although the details are species-specific (40). FtsZ treadmilling has been hypothesized to distribute PG synthesis and coordinate construction of the nascent endcaps by moving proteins around the division site (30); indeed, its dynamics have been confirmed to correlate with populations of moving PG enzyme complexes in *E. coli* (41) and *B. subtilis* (42), and in *C. crescentus* the FtsZ-binding partner FzlA links it to PG synthesis (43). In *B. subtilis*, FtsZ treadmilling is essential to mediate condensation of diffuse FtsZ filaments into a dense Z-ring, and to initiate constriction (44). Numerous lines of evidence suggest that FtsZ’s intrinsic capacity for polymerization provides the capability for Z-ring assembly, whereas its intrinsic GTPase activity is responsible for treadmilling dynamics, independent of other proteins or the cell cycle (45).

We propose that *Q* in our model may be identified with FtsZ, with the threshold value of 8 being the amount required for constriction initiation, and the time delay *T* the interval between constriction initiation and cell division. To support this proposition, we consider the supporting evidence first for *E. coli* then for *C. crescentus*. In *E. coli*, FtsZ is produced at a constant rate per unit volume, with cells accumulating FtsZ molecules constitutively to maintain a constant concentration of FtsZ (46), and constriction initiation coinciding with maximal Z-ring intensity (47), followed by rapid proteolytic degradation at the end of division (48). These observations are consistent with the picture in which FtsZ is produced at a rate proportionate to cell volume (represented by the effective stochastic Hinshelwood cycle variable *X*), with *k*_*Q*_ depending on condition-specific factors (35). Furthermore, a recent large-scale phenotyping study of *E. coli* across a range of nutrient conditions and perturbations observed that FtsZ is required for constriction initiation, which occurs after a constant mean cell length has been added, and that division follows constriction initiation with a constant time delay (49), consistent with the requirements of our model.

In the asymmetrically dividing *C. crescentus*, the picture is more complicated, as FtsZ levels are regulated in a cell cycle-dependent manner (50), with synthesis beginning slightly before swarmer cells differentiate into stalked cells and concentration reaching a maximum at the beginning of cell division before a precipitous drop (51). Transcription of *ftsZ* in swarmer and pre-divisional cells is repressed by the master cell cycle regulator CtrA (52), with transcription rates of *ftsZ* in stalked cells modulated by additional factors such as nitrogen and carbon availability (53). Although FtsZ is stable in the daughter stalked cell following cell division, it is cleared from the daughter swarmer cell (52, 54) via a regulated proteolysis that appears intrinsic to the asymmetric cell division of *C. crescentus* (55). Despite the complexity of this picture, we note that our model as written applies only to stalked daughter cells, in which case FtsZ dynamics satisfy the general requirements. Interestingly, in slow-growing *E. coli*, FtsZ synthesis displays a cell cycle dependence similar to that observed in *C. crescentus* swarmer cells (48). A future extension of our framework to incorporate these similar additional layers of control could yield insights into their implications for cell size homeostasis.

Alternatively, we may connect the identities in our model directly to the growth of cell surface, a complex process involving synthesis of PG precursors in the cytosol followed by final PG units at the cell surface (56). PG precursor synthesis is expected to occur in a cell cycle-independent manner and has been proposed as a regulator between growth and division, with accumulation of excess PG precursor material serving as a potential checkpoint for constriction initiation (57). Intriguingly, a mechanical homeostatic mechanism has been proposed to balance surface PG synthesis with overall cell growth rate (58). In stalked *C. crescentus* cells, PG synthesis occurs in an FtsZ-dependent manner, leading to medial elongation prior to Z-ring formation and predominantly midcell constriction thereafter (33, 59). In *E. coli*, preseptal synthesis is less important to cell elongation (60), although a similar “competition” between elongation and constriction for PG synthesis has been reported (61). In this picture, we connect *Q* to a component of PG synthesis (cytosolic PG precursors or surface PG subunits), with Θ and *T* remaining the initiation of constriction and the interval between constriction initiation and cell division, respectively. FtsZ dynamics then play an essential role in controlling the onset of constriction, with the thresholded species *Q* connecting cell elongation to the cell division machinery via an unknown mechanism.

### Stochastic behavior of T

We now consider stochasticity in the intergenerational evolution of size arising from the the division process duration *T* . Inspired by the phenomenon of FtsZ treadmilling, in which it has been observed that divisome proteins follow treadmilling filaments by a diffusion-and-capture mechanism as the process of cell wall constriction occurs (62), we speculate that the constriction-controlled division process may be approximately modeled as onedimensional drift combined with diffusion, where the traveling entity must traverse a certain distance to complete the process of division. (Perhaps the division machinery must move with the treadmilling FtsZ filaments around the Z-ring a certain number of times). In that scenario, *T* can be modeled as a first passage time (FPT) for one-dimensional drift with diffusion, whose distribution is an inverse Gaussian (14, 63):

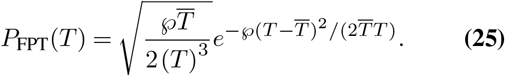

Herein, *𝓅* is the dimensionless Péclet number characterizing the drift-diffusion process and 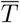 is the mean FPT. (If the process involves drift velocity *v*, diffusion constant *D*, and traversal length *ℓ*, then the Péclet number *𝓅* = *ℓ v/D*.) The experimentally determined *T* -distribution along with its fit with an inverse Gaussian is shown in Appendix Fig. 1. From the fit, we deduce a rough estimate of the Péclet n<u>u</u>mber *𝓅 ≈* 14 governing the underlying process (mean FPT 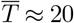 min).

## Concluding remarks

Observations of single *C. crescentus* cells with genetically and chemically perturbed constriction rates have demonstrated a role for constriction rate in size control and homeostasis (64). Future studies applying our framework to observations of single cells with perturbed constriction rates (i.e., modified *T* distributions), will yield further insights into the mechanism of stochastic intergenerational homeostasis under diverse conditions.

The mechanistic scheme we have proposed here displays the common property of control systems that the set of parameter values that give rise to the same emergent dynamics (constrained by Eqs. (18) and (19)) though infinitely large is vanishingly thin compared to the set of all possible parameter values. This allows for large situation-dependent variation in internal parameters while conserving the intergenerational size dynamics. In addition, preserving the constraints on certain protocols in our model allows for deconstraining other aspects of cellular processes while robustly maintaining cell size homeostasis. As growth conditions and the quality of available nutrients change, different underlying molecular circuitry may be involved in condition specific ways. Thus, the net effect may be to alter the underlying stochastic Hinshelwood cycle and hence the rates *k*_*X*_, *k*_*Q*_ and 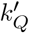, and also *T* . However, these alterations do not result in a breakdown of homeostasis since the homeostasis mechanism is indifferent to specific values of the rates of production of *Q* and *X*, provided the basic feature of initiation of division upon *Q* crossing the threshold is retained across conditions.

## ACKNOWLEDGMENTS

We thank Purdue University Startup funds, Purdue Research Foundation, the Purdue College of Science Dean’s Special Fund, and the Showalter Trust for financial support. K.J.and S.I.-B. acknowledge support from the Ross-Lynn Fellowship award and the Bilsland Dissertation Fellowship award. We are grateful to Iyer-Biswas group members for useful discussions. The datasets for constant growth conditions at 34°C utilized in this manuscript are published in (6).

## AUTHOR CONTRIBUTIONS

K.J., R.R.B., and S.I.-B. conceived of and designed the research; K.J. developed the theoretical framework under the guidance of S.I.-B.; K.J., R.R.B. and S.I.-B. performed analytic calculations; K.J. performed data analyses and simulations under the guidance of R.R.B. and S.I.-B.; C.S.W. made connections to molecular biological contexts with input from K.J. and S.I.-B.; K.J., C.S.W., R.R.B., and S.I.-B. wrote the paper; S.I.-B. supervised the research.

